# Experience-dependent alteration of mnemonic representation in early visual cortex and intraparietal sulcus

**DOI:** 10.1101/2020.10.20.347179

**Authors:** Ke Jia, Ya Li, Mengyuan Gong, Hui Huang, Yonghui Wang, Sheng Li

## Abstract

The ability to discriminate between stimuli relies on a chain of neural operations associated with perception, memory and decision-making. Accumulating studies show learning-dependent plasticity in perception or decision-making, yet whether perceptual learning modifies mnemonic processing remains unclear. Here, we trained participants on an orientation discrimination task, while using fMRI and TMS to separately examine training-induced changes in working memory (WM) representation. Although fMRI decoding revealed orientation-specific neural patterns during delay period in early visual cortex (V1) before, but not after, training, neurodisruption of V1 during delay period led to behavioral deficit in both phases. In contrast, both fMRI decoding and disruptive effect of TMS showed that intraparietal sulcus (IPS) represent WM content after, but not before, training. These results suggest that sensory engagement for WM is relatively independent of training but the coding format may be altered, whereas the involvement of parietal area in WM depends on training.

## Introduction

The ability to differentiate between similar features is essential for visual recognition in complex environment. For instance, the predators must learn to discriminate the prey items from surroundings to ensure survival. Learning and experience are known to improve the discrimination ability even in adulthood by re-organizing the brain functions and connections (Dosher & Lu, 2017; Gilbert et al., 2009; Hooks & Chen, 2020; Kourtzi & DiCarlo, 2006; Sagi & Tanne, 1994; Watanabe & Sasaki, 2015). Previous studies have focused on how training alters perceptual encoding of the stimuli (Chen et al., 2015; Furmanski et al., 2004; Jehee et al., 2012; Schoups et al., 2001; Schwartz et al., 2002; Yan et al., 2014; Yang & Maunsell, 2004; Yotsumoto et al., 2008) or the decision-making process (Dosher & Lu, 2017; Kahnt et al., 2011; Kuai et al., 2013; Law & Gold, 2008). However, mnemonic processing also matters for discrimination judgments where the to-be-compared stimuli are often sequentially presented. In these tasks, participants are required to encode a sample item and hold it in working memory (WM) for later comparison with a test item. Yet whether and how training on these tasks modifies the mnemonic processing of stimuli remain largely unclear.

The view that perceptual learning may change mnemonic processing of stimuli received support from findings of the relationship between WM and discrimination ability (Brady et al., 2013; Cornette et al., 2001; Ester et al., 2014; Zhang et al., 2016). In particular, variability of neuronal activity during WM retention is proposed as a potential indicator of the discrimination performance (Hussar & Pasternak, 2010; Qi & Constantinidis, 2015), the amount of information carried by the patterns activity during WM delay correlates with the mnemonic precision (Ester et al., 2013) and performance changes as a function of WM load (Emrich et al., 2013). These findings point to the assumption that learning-dependent improvement of discriminability may be accompanied by modified WM representation of the stimuli. It has been established that multiple levels of cortical areas are recruited for representing WM information (Christophel et al., 2017; D’Esposito, 2007; Dotson et al., 2018). In particular, intraparietal sulcus (IPS) area is identified as a candidate region for mnemonic processing of the stimuli (Bettencourt & Xu, 2016; Lorenc et al., 2018; Song & Jiang, 2006; Weber et al., 2016), while the sensory recruitment account of WM suggests that early visual area (V1) is also engaged for temporary maintenance of WM content (Ester et al., 2009; Harrison & Tong, 2009; Pasternak & Greenlee, 2005; Serences et al., 2009). Representing WM content at multiple areas could play complementary roles such that sensory areas encode precise sensory information and higher-order areas provide abstract and robust representation (Christophel et al., 2017; D’Esposito, 2007). Here, we particularly focus on these two areas to examine learning-dependent alterations of WM representation.

To this end, we trained participants on a two-interval forced-choice (2IFC) orientation discrimination task that required temporary maintenance of the sample stimulus during a delay period. In Experiment 1, we combined functional magnetic resonance imaging (fMRI) with multivariate pattern analysis (MVPA) to examine how learning moulds WM representation of the stimuli in V1 and IPS. We found orientation-specific patterns during WM delay in V1 before, but not after, training. In contrast to the orientation-specific patterns in IPS after, but not before, training. To further examine the causal role of these two areas in mnemonic processing along with training, in Experiment 2, we used online repetitive transcranial magnetic stimulation (rTMS) to disrupt V1 processing during delay period. We found that TMS over V1 during delay period led to behavioral deficit both before and after training, whereas TMS over IPS impaired performance after, but not before, training. These findings suggest differential effects of perceptual learning on mnemonic processing at different cortical levels. Sensory area engaged for WM representation is relatively independent of training, but the coding format may be altered such that it becomes insensitive to fMRI decoding. Higher-order parietal area is involved in WM depending on training, which may contribute to robust maintenance of feature information.

## Materials and Methods

### Experiment 1: fMRI

#### Participants

Sixteen participants (9 females; age range: 18 - 26 years) took part in this study. We determined the sample size that is comparable to those reported in previous work on perceptual learning (Zhang et al., 2010) or fMRI decoding of WM content using discrimination tasks (Ester et al., 2009; Gosseries et al., 2018; Lawrence et al., 2018). All participants had normal or corrected-to-normal vision, and reported being right-handed. They were naïve to the aim of the study and received payment upon completion of the experiment. All participants gave written informed consent and the study protocol was approved by the local ethics committee.

#### Stimulus and Apparatus

We presented Gabor patches (Gaussian windowed sinusoidal gratings) in either upper-left or lower-right visual field with an eccentricity of 6.5° against a gray background (~35 cd/m^2^). The Gabor stimuli of random phase had a fixed diameter of 4°, contrast of 0.8, spatial frequency of 1.5 cycle/degree. The angle of Gabor stimuli was tilted clockwise or counterclockwise from the base orientations (55° or 145°).

The stimuli were generated using Psychtoolbox 3.0 (Brainard, 1997; Pelli, 1997) for MATLAB (MathWorks, MA, USA). In the behavioral lab, the stimuli were presented on a Dell cathode ray tube monitor (CRT) with the size of 40 × 30 cm^2^, resolution of 1024 × 768 and a refresh rate of 60 Hz. Gamma correction was applied to the monitor. We used a chin-rest to stabilize participants’ head position and maintain the viewing distance at 90 cm. Participants were asked to make responses using a keyboard. Inside the MRI scanner, the stimuli were back-projected onto a translucent screen located inside the scanner bore (resolution, 1024 × 768; refresh rate, 60 Hz). Participants viewed the stimuli at a distance of 90 cm through a mirror placed above their eyes. A MRI-compatible response box was used for making responses.

#### Experimental procedure and tasks

All participants completed four phases in this experiment, each phase consisted of multiple sessions (Fig 1A): (1) a 2-day pretest, (2) a 6-day training, (3) a 2-day posttest I; and (4) a 2-day posttest II. Posttest I and II phases were separated for around ten days to assess the stability of training effect. Each test phase comprised of a behavioral test session (1^st^ day) and a scanning session (2^nd^ day) on two consecutive days.

**Fig 1.**
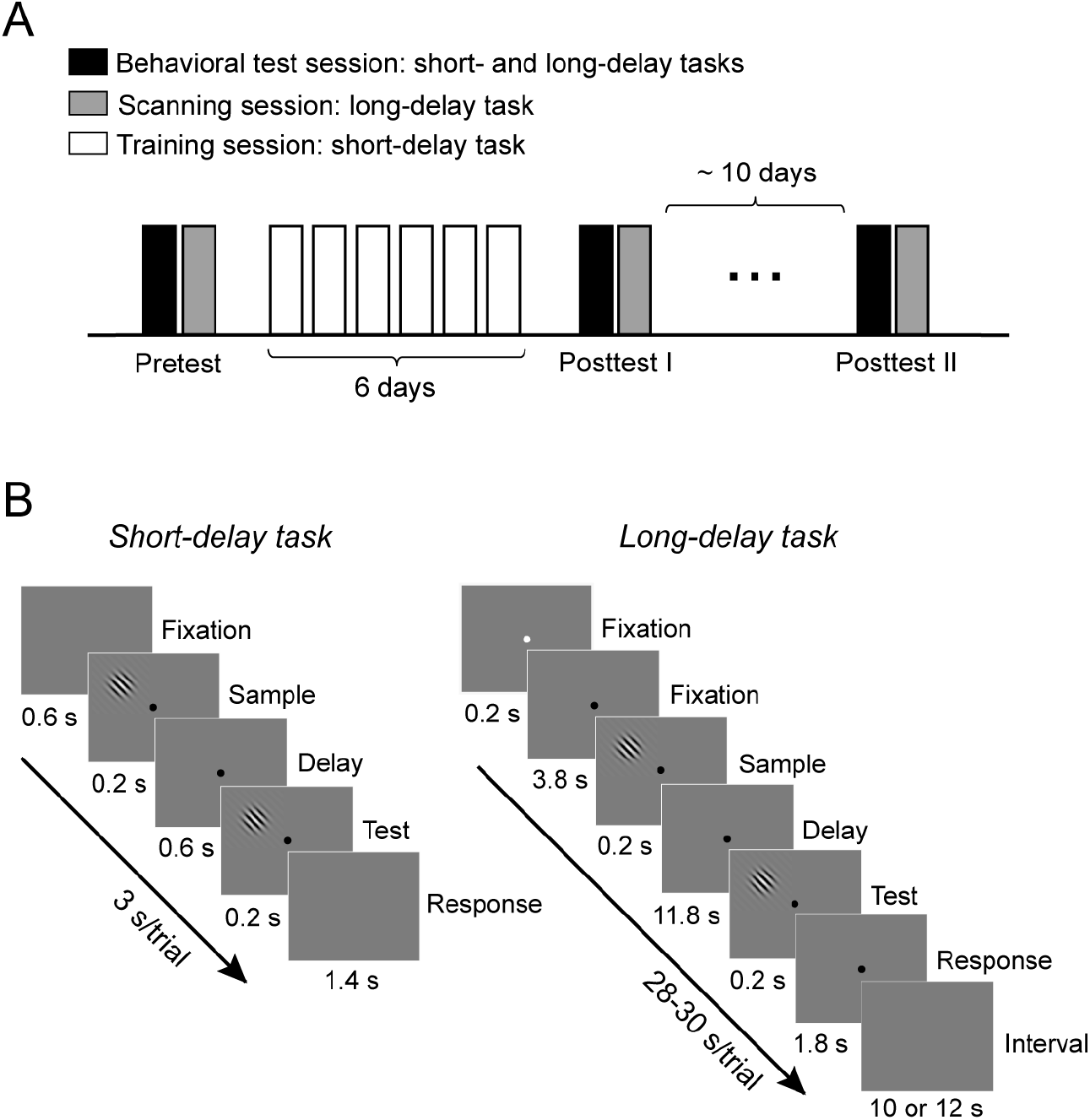
Experimental procedure and tasks. (A) Participants completed four phases in the experiment, including (1) a 2-day pretest, (2) a 6-day training, (3) a 2-day posttest I and (4) a 2-day posttest II. Each test phase comprised of a behavioral and a scan session, as separately denoted by black and gray bars. The first session of each test phase (behavioral session) included both short- and long-delay tasks. The second session of each phase (scanning session) included the long-delay task to address neural activity during WM delay. Training phase used short-delay task only. (B) Trial sequence of a short-delay (*Left*) and long-delay (*Right*) orientation discrimination tasks. Participants viewed two sequentially presented stimuli and reported whether the test stimulus was tilted clockwise or counterclockwise relative to the sample stimulus in both tasks.

##### Behavioral tasks

We used a 2IFC, orientation discrimination task throughout the experiment. Two types of tasks – a short-delay and a long-delay task – varying in the length of delay period between the stimuli were included (Fig 1B and 1C). Similar to the conventional learning regimen, we used a short-delay of 0.6 s to measure behavioral performance during the training and behavioral tests. To isolate memory-specific activity from the fMRI signal (Harrison & Tong, 2009; Serences et al., 2009), we used a long-delay of 11.8 s during the behavioral tests and scanning sessions.

In the short-delay task, each trial began with a central fixation of a black dot shown for 0.6 s. In the long-delay task, each trial began with a central fixation dot that was white for 0.2 s and then turned into black for 3.8 s. The change of color was designed to remind participants of the trial onset. Participants were instructed to press a button once they saw the white dot. In both tasks, the sample and test Gabor stimuli were then sequentially presented for 0.2 s each, separated by a delay period (short-delay task: 0.6 s; long-delay task: 11.8 s). Participants were asked to report whether the test Gabor was tilted clockwise or counter-clockwise relative to the sample stimulus. A uniformly distributed jitter (±5°) was added to the base orientations (i.e., 55° or 145°) to encourage perceptual comparison between two Gabors in each trial, rather than direct retrieval of a constant stimulus template.

##### Staircase procedure

To equate task difficulty across different conditions throughout the experiment, we used adaptive staircase (3-down-1-up, 15 reversals, step size of 0.5°) that converges to 79.4% accuracy in the orientation discrimination tasks. We adjusted the angle difference between the sample and test stimuli independently for each condition. The threshold in each run was determined by the mean angle difference of the last 8 reversals.

##### Behavioral test sessions (1^st^ day of pretest, posttest I and II)

This session included both the short-delay and long-delay orientation discrimination tasks. The short-delay task consisted of four experimental conditions (2 stimulus orientations × 2 stimulus locations) in order to assess the effect of learning and the learning specificity for orientation and location. Participants started with 16 practice trials (4 trials per condition) using a fixed angle difference (10°) and then completed 12 staircase runs (3 staircases per condition in random order). For the first run of each condition, the starting angle difference was 8° with a step size of 0.5°. For the subsequent staircase runs, the starting value was the threshold of corresponding condition in the preceding run. Participants’ performance in each session was quantified using the averaged thresholds across three staircases for each condition.

To keep consistency with the trial sequence in the scanning session, the long-delay task consisted of two stimulus conditions (i.e., ~55° or ~145°) shown only at the trained location. Participants began with 20 practice trials (10 trials per condition, fixed angle difference: 10°) and then completed one run of randomly interleaved staircases (see Staircase procedure). The starting angle difference was 8° with a step size of 0.5°. We quantified the performance using the threshold for each condition. No feedback on correctness was provided in any of these test sessions.

##### Scanning sessions (2^nd^ day of the pretest, posttest I and II)

Participants completed 6 runs of a long-delay task (16 trials per run, 8 trials for each orientation in randomized order), each run began with an 8 s fixation. Trials were separated by a 10 s or 12 s interval to allow fMRI signals to return to baseline. We measured performance with staircase procedure. The starting value was the threshold inherited from the preceding behavioral session in the corresponding test phase. In addition to the discrimination task, each participant completed a retinotopic mapping scan (6 min 20 s), a localizer scan (5 min 36 s) and an anatomical image scan (see ROI definition for details). No feedback on the correctness was provided in the scanning sessions.

##### Training sessions

Participants were trained on an orientation discrimination task with Gabors presented at the same orientation and location throughout training. In each session, participants performed 16 runs of short-delay task. We measured performance with staircase procedure. For the first run of the first session, the starting angle difference was 8° with a step size of 0.5°. For the subsequent staircase runs, the starting value was the threshold from the preceding run. Training locations (i.e., upper-left or lower-right) and orientation (i.e., 55° or 145°) were counterbalanced across participants. In addition, we provided auditory feedback on incorrect trials.

##### Behavioral data analysis

To validate the training effect, we used paired t-test to compare the discrimination threshold between the first and last sessions of the training phase. To examine the effect of training on discrimination performance in the test phases, we calculated a mean percent improvement (MPI = (pretest threshold – posttest threshold)/pretest threshold × 100%) (Xiao et al., 2008), separately for each posttest phase. For the short-delay task, we applied a three-way repeated-measures ANOVA (2 stimulus orientations × 2 stimulus locations × 2 posttest phases) on MPI. For the long-delay task, we applied a two-way repeated-measures ANOVA (2 stimulus orientations × 2 posttest phases) on MPI due to the presence of stimulus solely at the trained location.

#### MRI data acquisition and preprocessing

Imaging data were acquired on a Siemens 3T Prisma scanner located at Peking University. All imaging data were acquired with a 20 channel head coil. For each participant, anatomical images were acquired using MPRAGE T1-weighted sequence (TR = 2530 ms, TE = 2.98 ms, FOV = 256 × 224 mm^2^, flip angle: 7°, resolution 0.5 × 0.5 × 1 mm^3^, number of slices: 192, slice thickness: 1 mm, slice orientation: sagittal). Functional scans were acquired using echo planar imaging (EPI) sequence (TR = 2000 ms, TE = 30 ms, FOV = 224 × 224 mm^2^, flip angle: 90°, matrix: 64 × 64, resolution 3.5 × 3.5 × 3.5 mm^3^, gap = 0.7 mm, number of slices: 33).

Each participant’s anatomical image was segmented into gray and white matter using FreeSurfer (http://surfer.nmr.mgh.harvard.edu/). We performed the cortical reconstruction of the segmented images in BrainVoyager QX software (Brain Innovation, Maastricht, The Netherlands). For the functional images, we discarded the first four volumes at the beginning of each run to ensure that the longitudinal magnetization reached steady state. The functional data was processed with slice-timing correction, head motion correction, temporal filtering (3 cycles) and removal of linear trends in BrainVoyager QX. Within each scanning session, the functional data was aligned to the first volume of the first run and co-registered to the anatomical image obtained in the same session. Between scanning sessions, all anatomical images were aligned to the participant’s own anatomical data acquired in their first session and transformed to the Talairach space. The functional data in the Talairach space were resampled into 3 × 3 × 3 mm resolution.

#### ROI definition and fMRI data analysis

##### Definition of V1

Participants viewed rotating wedges that created travelling waves of neural activity (Engel et al., 1997; Sereno et al., 1995). We identified V1 boundaries using standard phase-encoded method. In a separate localizer run, we mapped two location-specific areas in V1 corresponding to the stimulus locations from the orientation discrimination task (i.e., upper-left and lower-right). In each trial, a Gabor patch was presented at one of the locations for 2 s. The inter-trial interval was either 2 s or 4 s. The location was randomized across 32 trials. Participants were asked to detect a subtle change of orientation. For each participant and each functional localizer, we computed each voxel’s response using a General Linear Model (GLM) comprised of two regressions of the stimulus locations. We selected 40 voxels with top-ranked beta estimates (β) for each stimulus location, the exact number of voxels was determined by the minimal number of voxels across participants and V1 ROIs. This voxel selection regime controlled for potential biases in classification accuracy due to varying number of voxels across locations and participants.

##### Definition of IPS

We defined two parietal ROIs (i.e., left and right intraparietal sulcus, IPS) given their functional relevance to the retention of WM. After applying anatomical segmentation in FreeSurfer, we used the automated ROI labels from Destrieux atlas (Destrieux et al., 2010) to transform the identified IPS into Talariach space. For each participant, we conducted a GLM analysis that modelled the WM-related activity after the sample stimulus (i.e., delay period) and the baseline activity after the test stimulus (i.e., inter-trial interval). The resulting β estimates that indicated statistically significant increases of delay-period activity (*p* < 0.05) were used for voxel selection within anatomically-defined ROIs (Xu, 2007). In total, we selected 250 voxels with top-ranked β estimates in each hemisphere for further analysis, the exact number of voxels was determined by the minimal number of voxels across participants and ROIs.

##### Univariate analysis

We assessed whether training changes the overall BOLD response during WM delay. For each participant and each run, we first extracted z-normalized response amplitude of each voxel in the pre-defined ROIs (V1 and IPS). Then, we took the trial-averaging BOLD response between 0 – 26 s time locked to trial onset, separately for each experimental condition during the test phase (Figure 3). Because of our primary focus on the sustained activity during the delay period, we defined a window from 12 s to 14 s after the trial onset to minimize the influence of the sample or test stimulus-related activity. For each of the ROIs, we applied a two-way repeated-measures ANOVA (2 stimulus orientations × 3 test phases) on the delay activity to assess how training influenced WM-related activity.

##### Multivariate pattern analysis (MVPA)

We used the MVPA to decode the stimulus orientation during the delay period in V1 and IPS. For each participant and each test phase, we separately extracted z-normalized BOLD responses between 12 s to 14 s after the trial onset (i.e., 8 s to 10 s after the onset of the sample stimulus) in each trial that represented the delay period activity. By training the classifier to discriminate between two orientations using LIBSVM (http://www.csie.ntu.edu.tw/~cjlin/libsvm/), we calculated the classification accuracy with a leave-one-run-out validation scheme that divided the data set into training (5 runs) and testing data (1 run). This procedure was repeated for 6 times until each run was tested once (Kamitani & Tong, 2006). The classification accuracy was averaged across the folds, separately for each test phase. To examine whether orientation decoding reflected any residual sensory information, we applied MVPA on the neural activity between 24 s to 26 s after the trial onset (i.e. 8 s to 10 s after the onset of the test stimulus). Note that for V1, the ROIs were classified as contralateral versus ipsilateral locations corresponding to the stimulus locations in the visual field. For IPS without precise mapping of stimulus locations, we defined ROIs in the left and right hemisphere. To assess the statistical significance of the decoding accuracy in each ROI, we used one sample t-tests against the chance level (50%).

### Experiment 2: TMS

#### Participants

Twenty-three participants (12 females; age range: 19 – 26 years) were recruited for this study. Three of them did not participate TMS sessions after fMRI scanning due to the lack of elevated delay period activity in IPS (see ROI definition for details). The sample size was comparable to those reported in previous WM-related TMS studies (Zanto et al., 2014; Zokaei et al., 2014). All participants were neurologically intact, had normal or corrected-to-normal vision, and reported being right-handed. All participants gave the informed consent and the study protocol was approved by the local ethics committee.

#### Stimulus and Apparatus

Identical stimuli were used as that in Experiment 1. The stimuli were presented against a gray background (~15 cd/m^2^) on a CRT monitor (refresh rate: 60 Hz) for both behavioral and fMRI experiments. In the TMS lab, the stimuli were displayed on a gray background (~19 cd/m^2^) on a CRT monitor (refresh rate: 100 Hz).

#### Experimental procedure and tasks

The TMS experiment consisted of four phases: (1) a scan for defining ROIs (V1 and IPS) per participant, (2) a 2-day pretest, (3) a 6-day training, (4) a 2-day posttest. Pretest and posttest were completed one day before and after the training phase, respectively. The pre- and posttest phases consisted of a behavioral session (1^st^ day) and a TMS session (2^nd^ day).

##### Scanning session

To guide precise stimulation of the target regions in TMS sessions, each participant completed a V1 localizer scan (2 runs) and an IPS localizer scan (1 run), in addition to an anatomical image scan and a retinotopic mapping scan (see ROI definition for details).

##### Behavioral test sessions (1^st^ day of the pretest and posttest)

Participants performed a long delay orientation discrimination task that was similar to that used in Experiment 1. In brief, the total duration of each trial was fixed to 7 s. Each trial began with a central fixation of a black dot shown for 0.6 s. The sample and test Gabor stimuli were then sequentially presented for 0.2 s each, separated by a 4-s delay. A central fixation against the gray background followed until a response was made or wait until 1.5 s. A blank screen was presented in the remaining time as the inter-trial interval. Participants were asked to report whether the test Gabor was tilted clockwise or counter-clockwise relative to the sample stimulus. Notable changes were made to the task design for several practical concerns. First, we shortened the delay period from 11.8 s to 4 s during online TMS stimulation. The sluggish BOLD signals require a long delay to isolate WM-related activity, which is not necessary for assessing the effects of neurodisruption. Second, following our fMRI results of feature-specific information in left IPS, we specifically presented the stimulus at the lower-right visual field that corresponds to the left hemisphere. Third, we used one orientation (55°) as we aimed to compare the TMS effects on discrimination performance between test phases (i.e., pretest vs. posttest) and between stimulation sites (i.e., V1, IPS and vertex), rather than between two orientations (i.e., the trained and untrained orientations). Participants started with 40 practice trials (a fixed angle difference: 10°) and then completed 2 to 3 runs of the main task using the same staircase procedure, identical to that used in the behavioral test sessions of Experiment 1. No feedback on correctness was provided in these sessions except practice trials.

##### Training sessions

Participants were trained to discriminate the orientation around 55° presented at the lower-right visual field. On each session, they performed 16 staircase runs, using the same protocol as that in the training sessions of Experiment 1. We provided auditory feedback on incorrect trials.

##### TMS sessions (2^nd^ day of the pretest and posttest)

In an orientation discrimination task, we used a fixed angle difference determined by the threshold from the behavioral test session in corresponding phase for each participant. Participants started with 40 practice trials and then completed 3 runs (80 trials per run). In separate runs, the magnetic stimulation was delivered to one of the three stimulation sites (V1, IPS and vertex) during the delay period. The order of the stimulation site was counterbalanced across participants. No feedback on correctness was provided in the TMS sessions.

#### TMS and MRI parameters

##### MRI data acquisition and preprocessing

Imaging data were acquired on a 3T GE MEDICAL SYSTEMS scanner located at Peking University using an 8-channel head coil. For each participant, anatomical images were acquired using T1-weight sequence (TR = 6.656 ms, TE = 2.92 ms, FOV = 256 × 256 mm^2^, flip angle: 90°, resolution 1 × 1 × 1 mm^3^, number of slices: 192, slice thickness: 1 mm, slice orientation: sagittal). Functional scans were acquired using EPI sequence (TR = 2000 ms, TE = 30 ms, FOV = 224 × 224 mm^2^, flip angle: 90°, matrix: 64 × 64, resolution 3.5 × 3.5 × 3.5 mm^3^, gap = 0.7 mm, number of slices: 33). The same preprocessing procedure was applied to fMRI data as that in Experiment 1.

##### Definition of ROIs

For each participant, the functional data were aligned to the anatomical data in native space. We defined V1 using retinotopic mapping and standard phase-encoded method. Further, we applied a GLM on data from two localizer runs to estimate each voxel’s response in V1 (i.e., beta estimate, β), allowing us to define the exact stimulus location (β: lower-right > upper-left). Identical to our voxel selection approach in Experiment 1, we selected IPS voxels showing elevated delay period activity (β: delay period > inter-trial interval), while also locating within an anatomically defined IPS ROI. In addition, we included the vertex as a control site for sham TMS. Vertex was defined as a midpoint between inion and nasion that was equidistant from left and right intertrachial notches. The coil was centered at the vertex with its face rotated 90° away from the scalp during stimulation. Thus, no cortical stimulation should be received during sham TMS.

##### Repetitive TMS (rTMS) protocol

To investigate the cause role of sensory and parietal areas during WM retention in discrimination performance along with training, online rTMS was applied over V1 and IPS during the delay period. Online 10 Hz rTMS (5 pulses synchronized with 1500 ms after the offset of the sample stimulus) was delivered at each stimulation site with a fixed intensity of 60% of the stimulator’s maximum output for all participants (Chang et al., 2014; Mevorach et al., 2010). In particular, we included two target sites contralateral to the trained location (i.e., left V1 and IPS) and a control site (i.e., vertex) for sham TMS. rTMS pulses were delivered through a MagStim Super Rapid^2^ stimulator (The MagStim Company, UK) in combination with a 70-mm figure-of-eight coil. This TMS protocol was used to produce interference effect (Chang et al., 2014; Mevorach et al., 2010; Silvanto & Soto, 2012) and disrupt BOLD signal in stimulated area in a simultaneous fMRI-TMS study (Sack et al., 2007).

Using fMRI-guided Visor Navigation System (Visor2; Advanced Neuro Technology, Enschede, The Netherlands), we separately overlaid V1 and IPS on the anatomical MR image for each participant with their centroid serving as the target site. The center of the coil was placed tangentially over these sites and a mechanical arm was used to keep the coil steady on the scalp. During V1 stimulation, the coil was held with the handle pointing right and parallel to the ground. During IPS stimulation, the coil was held with the handle pointing away ~45° along the midline (Capotosto et al., 2012; Morgan et al., 2013). The coil position in different sites was chosen based on the literature (Janssen et al., 2015) and was in real-time monitored using Visor2 throughout each session.

#### Behavioral analysis

Behavioral performance was quantified using the discrimination accuracy and reaction time (RT). To validate the effect of training, we used paired t-test to compare the discrimination thresholds between the first and the last sessions of training phase. To unravel TMS specific effect, we separately compared the performance from the active TMS (V1 and IPS) to the sham TMS condition to rule out non-specific effects related to variations in general behavioral state (e.g., noise, vigilance). We applied two separate two-way repeated-measures ANOVAs (2 TMS conditions: active vs. sham × 2 test phases: pretest and posttest) on the discrimination accuracy and RT.

## Results

### Experiment 1: fMRI

#### Perceptual learning improves performance in short and long delay tasks

Perceptual learning improved participants’ discrimination performance, as revealed by the decreased threshold from the first session (Mean = 3.00°, SD = 0.81°) to the last session (Mean = 2.11°, SD = 0.51°) of the training phase (Fig 2A, paired t-test: t(15) = 6.40, *p* < 0.001, Cohen’s d = 1.60).

**Fig 2.**
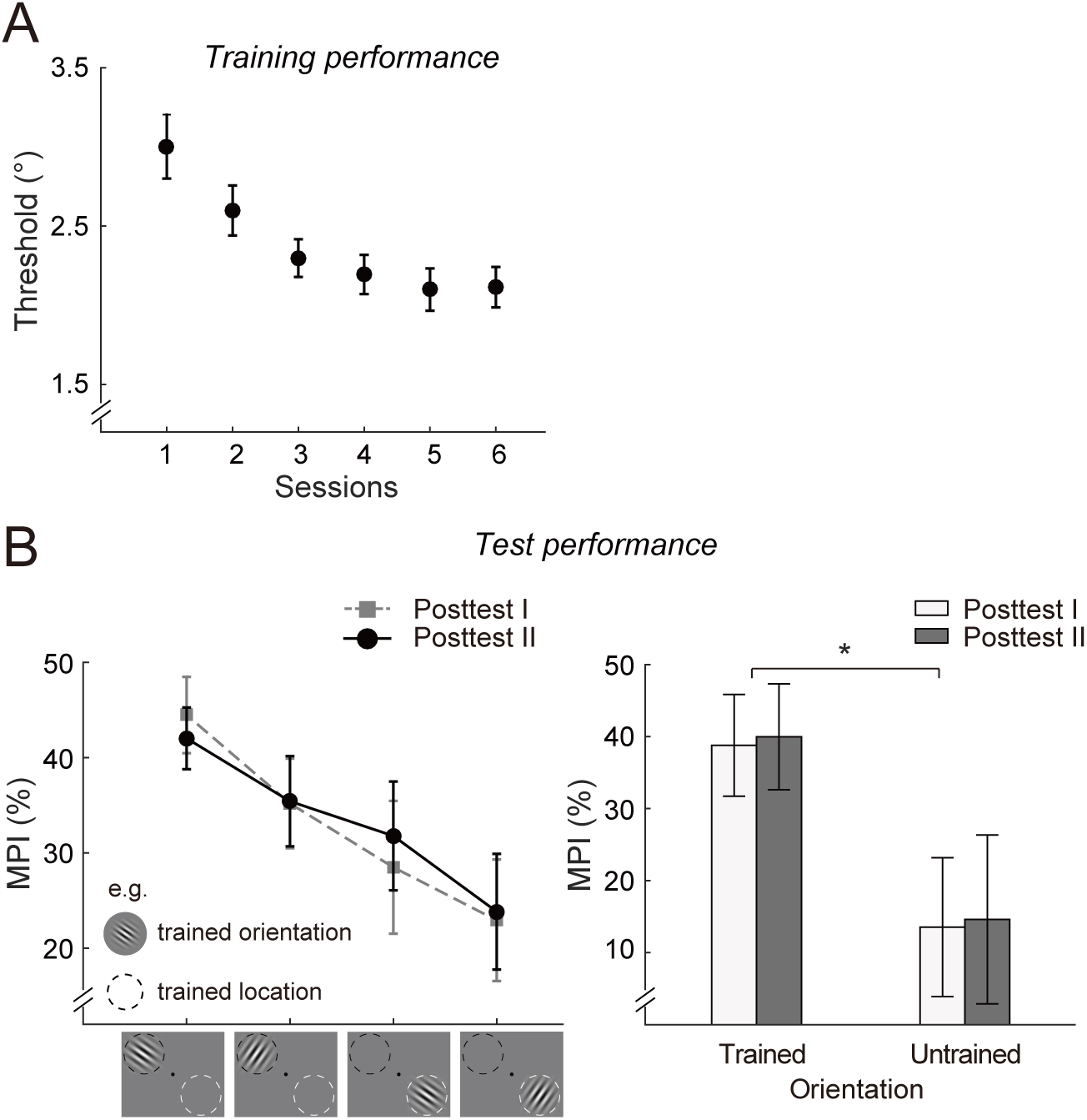
Behavioral results of Experiment 1. (A) Participants’ discrimination threshold over training sessions. (B) Participants’ performance during the posttest phases. (*Left*) Mean percent improvement (MPI) in the short-delay task during the posttest phase I and II. As shown in an example condition (e.g., training 145° at the up-left location), MPI was highest for the trained orientation (145°) at the trained location (black dashed circle). (*Right*) MPI in the long-delay task during the posttest phase I and II. Error bar represents standard error of the mean. The symbols indicate significant difference (\**p* < 0.05).

To assess the specificity of learning effect, we applied a three-way repeated-measures ANOVA (2 stimulus orientations × 2 stimulus locations × 2 posttest phases) on MPI in the short-delay task (Fig 2B, left panel). The results showed a main effect of stimulus orientation (F(1,15) = 6.06, *p* = 0.026, 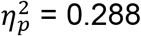) and stimulus location (F(1,15) = 12.51, *p* = 0.003, 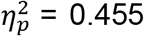), indicating more training-induced improvement for the trained versus untrained orientation, as well as training-induced improvement for the trained versus untrained location. None of the other main effects or interaction effects reached significance level (*ps* > 0.35). Importantly, when applying a two-way ANOVA (2 stimulus orientations × 2 posttest phases) on MPI in the long-delay task (Fig 2B, right panel), we also observed a significant main effect of the stimulus orientation (F(1,15) = 4.80, *p* = 0.045, 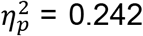), suggesting that training on the short-delay task improved discrimination performance in the long-delay task. None of the other effects reached significance level (*ps* > 0.80).

#### Perceptual learning does not change BOLD amplitudes in V1 and IPS

To examine whether perceptual learning changes the response amplitude in V1 and IPS during the delay period, we used event-related analysis that compared BOLD response between the trained and untrained orientations. Fig 3 showed an example of the averaged temporal dynamics of BOLD responses in V1 (i.e., contralateral and ipsilateral locations) and IPS (i.e., left and right IPS) from posttest I. The delay period activity almost dropped to baseline level in V1, whereas an elevated delay period activity was observed in IPS. This elevated delay period activity was generally assumed to reflect WM retention (D’Esposito & Postle, 2015; Sreenivasan & D’Esposito, 2019). A two-way repeated-measures ANOVA (2 stimulus orientations × 3 test phases) revealed no significant effects on the delay activity in either V1 (*ps* > 0.31 for all comparisons) or IPS (*ps* > 0.12 for all comparisons). These results suggest that training did not alter overall BOLD response during WM delay in sensory and higher-order areas, in line with prior studies (Lorenc et al., 2018).

**Fig 3.**
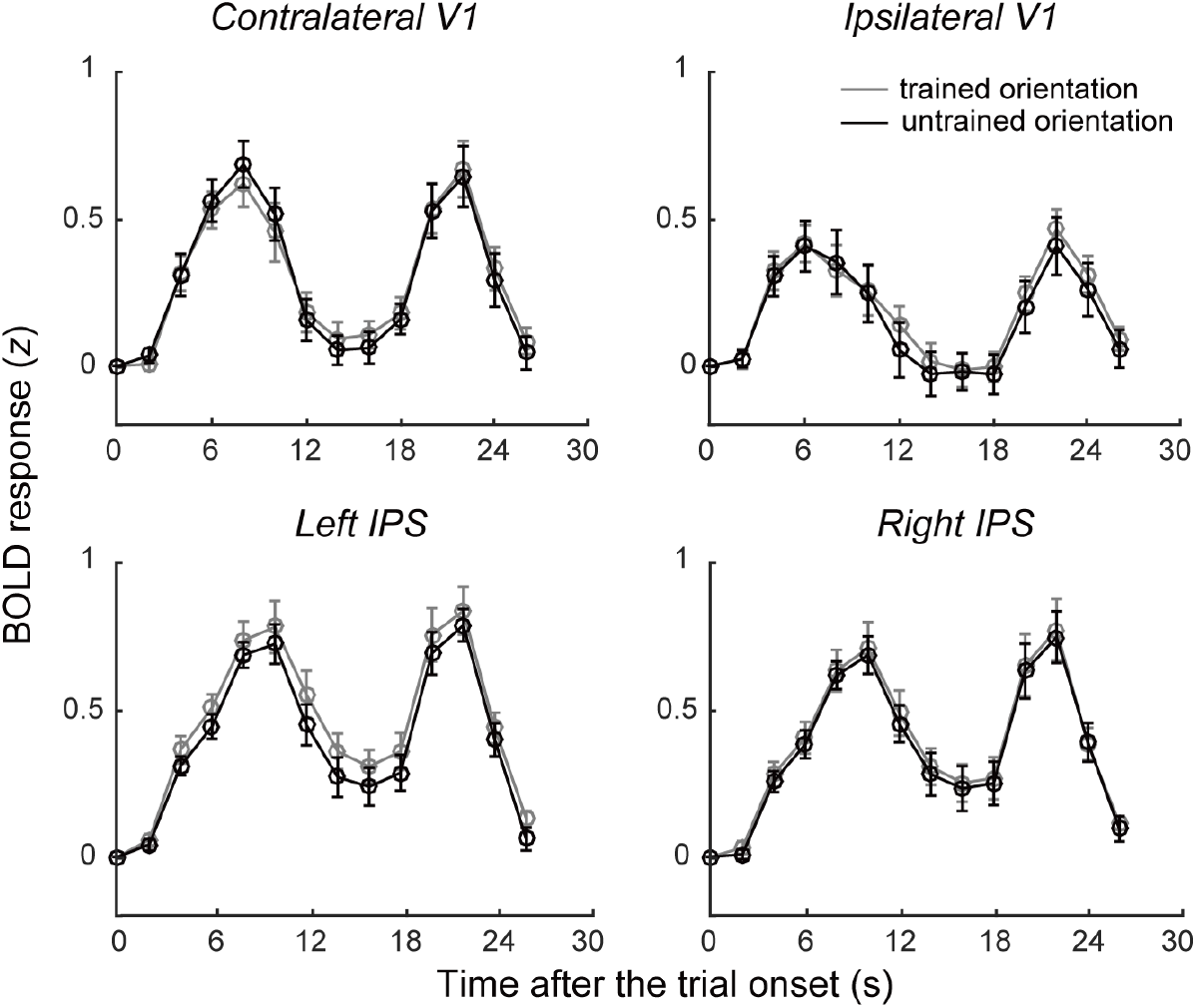
Time course of z-transformed BOLD activity in V1 and IPS in post-test I. First row represents contralateral and ipsilateral activity in V1. The second row represents activity from left and right IPS. Each subplot shows the time course of BOLD response for the trained and untrained orientations. Error bar represents standard error of the mean.

#### Discrimination training alters feature-specific WM representation in V1 and IPS

We next examined whether the feature-specific information was contained in the distributed pattern activity during delay period and how training modulated such representation. Using MVPA (Kamitani & Tong, 2006), we decoded the stimulus orientation in V1 during the delay period, separately for each test phase (Fig 4, *Left*). Repeated-measures ANOVAs on the classification accuracy in V1 revealed a main effect of test phases (pretest, posttest I and II) in the contralateral (F(2,30) = 5.72, *p* = 0.008, 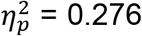) and the ipsilateral ROIs (F(2,30) = 5.63, *p* = 0.008, 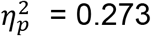). The decoding accuracies were above-chance in both ROIs before training (contralateral ROI: t(15) = 4.03, *p* = 0.001; ipsilateral ROI: t(15) = 3.63, *p* = 0.002, one-sample t-test). In contrast, classification accuracy dropped to chance-level after training for both contralateral and ipsilateral ROIs (*ps* > 0.11). To ensure that selected time window for MVPA reflected WM retention and was not spuriously induced by residual effect of sensory processing, we applied the same analysis to the data from inter-trial interval, a period following the test stimulus (i.e., 8 s to 10 s after the onset of the test stimulus). This period presumably contained comparable sensory information to that during the WM delay following the sample stimulus (i.e., 8 s to 10 s after the onset of the sample stimulus) but without the demand of WM maintenance. The results showed chance level classification accuracy after the test stimulus (*ps* > 0.12 for all comparisons), thus guarding against the possibility that V1 decoding of feature during WM delay was due to residual sensory processing.

**Fig 4.**
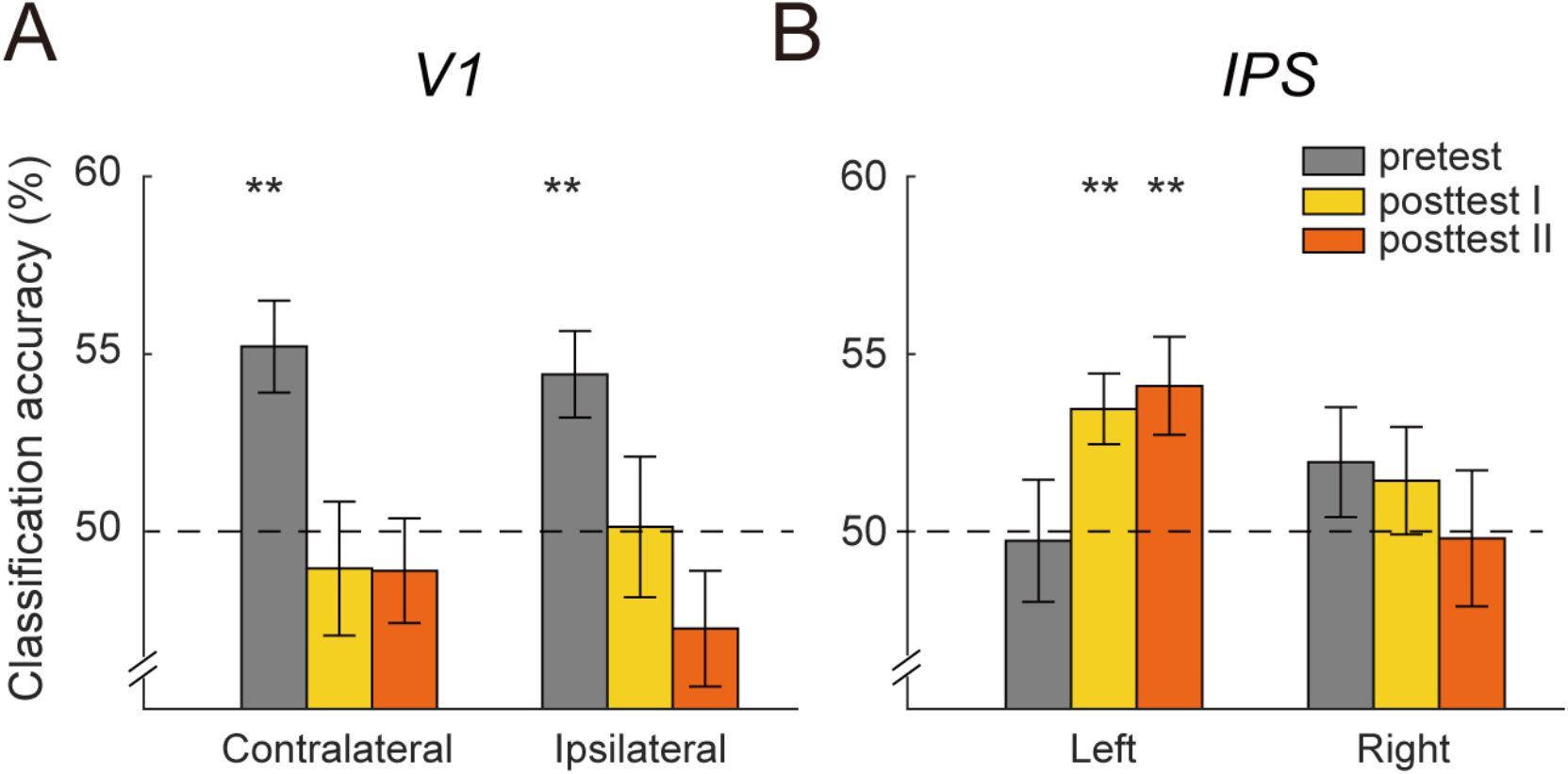
MVPA results of Experiment 1. (A) Orientation decoding in V1 and (B) IPS across three test sessions. The black dashed line represents the chance level (50%). Error bar represents standard error of the mean. The symbols indicate the significance level (\**p* < 0.05, \*\**p* < 0.01).

To address whether high-order areas related to WM processing contained feature-specific information and how training influenced such representation, we trained the classifier to distinguish between two orientations in bilateral IPS (Fig 4, *Right*). Repeated-measures ANOVAs revealed a main effect of test phase (pretest, posttest I and II) in left IPS (F(2,30) = 3.56, *p* = 0.041, 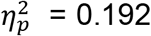), but not in right IPS (F(2,30) = 0.39, *p* = 0.684). In particular, we observed above-chance accuracies in left IPS after training (posttest I: t(15) = 3.46, *p* = 0.003; posttest II: t(15) = 2.97, *p* = 0.009), but not before training (pretest: t(15) = −0.15, *p* = 0.881). We did not observe reliable decoding in any of the test phases for right IPS (*ps* > 0.23). This left lateralization was probably anticipated, as the neural codes in left IPS was suggested to be more tightly associated with WM precision (Weber et al., 2016). To summarize, these multivariate fMRI results demonstrate distinct profiles of learning-dependent changes in classification performance between left IPS and V1.

### Experiment 2: TMS

#### Replication of the learning effect on behavior in Experiment 1

Participants were trained on a short-delay orientation discrimination task and showed decreased threshold over the training sessions (first session: Mean = 2.99°, SD = 0.93°; last session: Mean = 2.22°, SD = 0.51°; paired t-test: t(19) = 5.15, *p* < 0.001, Cohen’s d = 1.15). Similar learning effect was also observed in the long-delay task (pretest: Mean = 3.46°, SD = 1.11°; posttest: Mean = 2.53°, SD = 0.55°; paired t-test: t(19) = 3.62, *p* = 0.002, Cohen’s d = 0.81). These results replicated the behavioral effects observed in Experiment 1, showing that training enhanced discrimination performance in the long-delay task.

#### Discrimination training differentially impairs behavior with V1 and IPS stimulation

While our multivariate analyses in Experiment 1 suggest that perceptual learning changes the engagement of V1 and IPS during WM retention, we further took advantage of TMS to infer causal relation between neural and behavior and examine how such relation was altered after training.

The results from Experiment 1 showed reliable orientation decoding in V1 during WM delay before, but not after, training. These results provide a seeming account that V1 became unnecessary for WM after training. We thus disrupted V1 activity during WM delay and compared the change of performance to the sham condition (Fig 5A). A two-way repeated-measures ANOVA (2 test phases: pretest vs. posttest × 2 stimulation sites: V1 vs. vertex) on discrimination accuracy revealed a main effect of the stimulation site (F(1,19) = 11.05, *p* = 0.004, 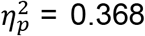), without a two-factor interaction or main effect of test phase (*ps* > 0.263). These results confirmed the validity of TMS-specific effects on V1 and further suggest the necessary role of V1 for WM retention after training.

**Fig 5.**
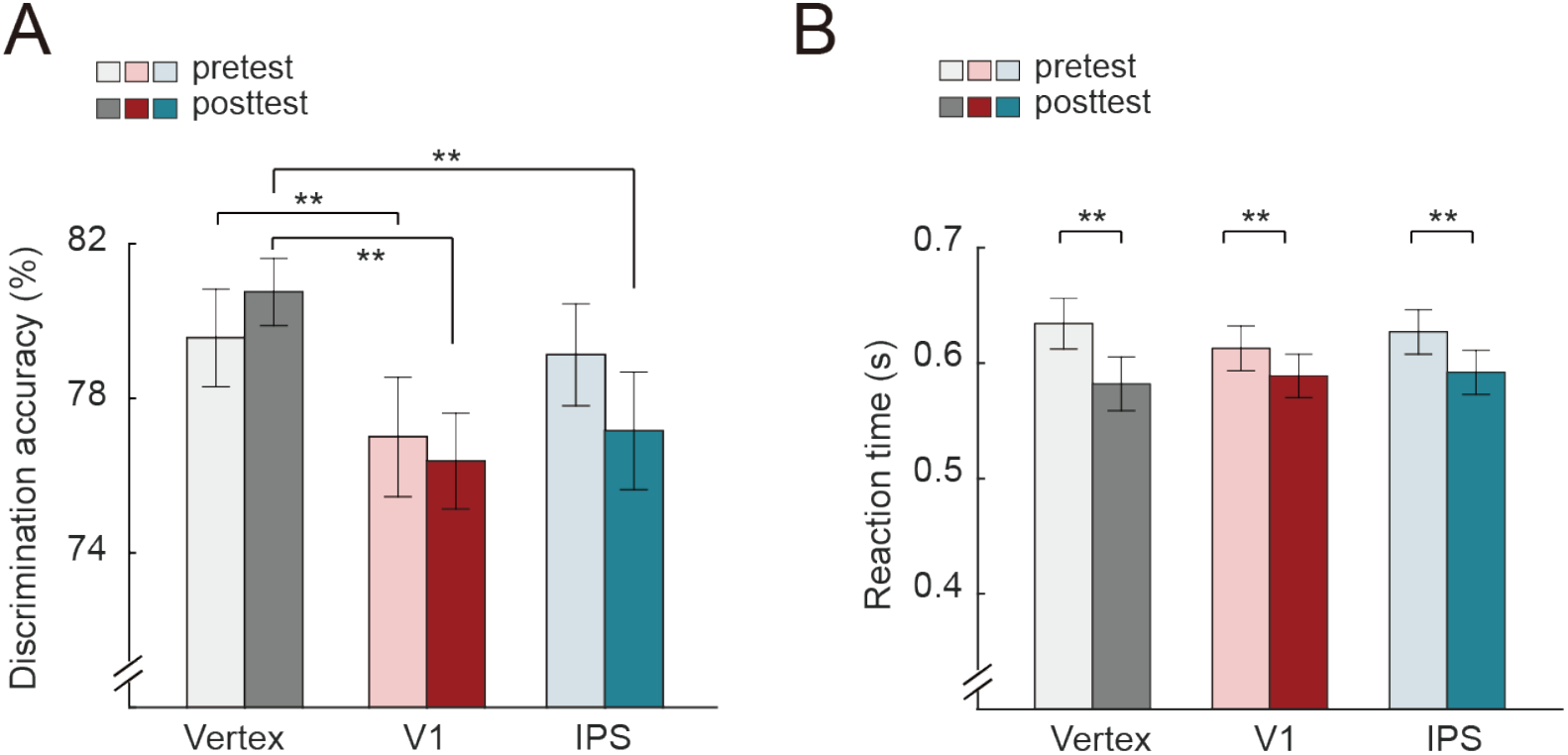
TMS results of Experiment 2. (A) Discrimination accuracy at the pretest and posttest phases. From left to right, each pair of bars corresponds to behavioral performance before and after training across different TMS conditions (vertex, V1 and IPS). (B) RT at the pretest and posttest phases. Error bar represents standard error of the mean. The symbols indicate the significance level (\**p* < 0.05, \*\**p* < 0.01).

On the contrary, the fMRI results showed reliable orientation decoding in IPS during WM delay after, but not before, training. These results predicted a learning-dependent involvement of IPS for WM maintenance. With this rationale, we disrupted IPS activity during WM delay and compared the change of performance to that in the sham condition (Fig 5A). A two-way repeated-measures ANOVA (2 test phases: pretest vs. posttest × 2 stimulation sites: IPS vs. vertex) on discrimination accuracy revealed a main effect of stimulation sites (F(1,19) = 8.38, *p* = 0.009, 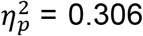). Importantly, we observed a two-factor interaction effect (F(1,19) = 4.56, *p* = 0.046, 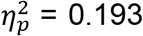). Simple effect analyses showed that stimulating left IPS significantly impaired performance at the posttest phase (IPS vs. vertex: t(19) = −3.29, *p* = 0.002, one-tailed), but not at the pretest phase (IPS vs. vertex: t(19) = −0.47, *p* = 0.322, one-tailed). These results converged with fMRI findings and jointly suggest that perceptual learning enhanced the involvement of IPS in WM maintenance.

To rule out the alternative possibility that the disruptive effect of TMS on the accuracy reflected the speed-accuracy tradeoff, we performed the same ANOVA tests on RT (Fig 5B). The results showed a main effect of test phase between V1 and vertex, and between IPS and vertex (*ps* < 0.001), suggesting an effect due to practice. Importantly, there were no interaction effects between V1 and vertex, or between IPS and vertex (*ps* > 0.15), eliminating the account of speed-accuracy tradeoff.

## Discussion

In the present study, we provide evidence that training alters mnemonic representation of simple visual features (i.e., orientation) in a discrimination task. We focused on V1 and IPS that have been associated with WM retention for visual features (Bettencourt & Xu, 2016; Weber et al., 2016; Harrison & Tong, 2009; Serences et al., 2009). In particular, combining fMRI and MVPA techniques (Experiment 1), we found orientation-specific information during WM delay that was decodable in V1 before, but not after, training; in contrast to decodable WM information in IPS after, but not before, training. Using online rTMS to disrupt WM delay-period activity in these two areas (Experiment 2), we found that neurodisruption of V1 led to behavioral deficits both before and after training, whereas neurodisruption of IPS led to selective deficit only after training. These findings thus point to learning-related changes in mnemonic representation of visual features that varied at different cortical levels (i.e., V1 vs. IPS), complementing prior studies that mainly addressed alterations of sensory and decision-making processes.

Previous neurophysiological and neuroimaging studies have shown training-induced changes in visual cortex that presumably occurred at an early stage of encoding processes (Chen et al., 2015; Furmanski et al., 2004; Jehee et al., 2012; Schoups et al., 2001; Schwartz et al., 2002; Yan et al., 2014; Yang & Maunsell, 2004; Yotsumoto et al., 2008). Here, we examined the contribution of visual cortex to mnemonic representation. Converging pre-training results from fMRI decoding and TMS provided direct support for the theoretical hypothesis of “sensory recruitment of WM” (Harrison & Tong, 2009; Pasternak & Greenlee, 2005; Serences et al., 2009). Extending beyond these findings, we further tested the effects of training on the sensory engagement for WM. Unexpectedly, while the post-training TMS results suggests a learning-independent mechanism of V1 during WM maintenance, we did not find decodable WM information in V1. Of note, the null result in V1 decoding after training was paired with positive results of V1 decoding before training, using the same sets of voxels and analytical approach. It is thus unlikely that this null effect reflected lack of sensitivity in voxel selection for decoding. Therefore, we speculate that training moulds the coding format in this region that is insensitive to decoding.

Although our observation of learning-independent behavioral impairment after V1 stimulation seems at odds with the inability to decode WM content in V1 specifically after training, such discrepancy may be reconciled by a time-varying attentional modulation on sensory processing given the close relationship between WM and attention (Awh & Jonides, 2001; Gazzaley & Nobre, 2012). Itthipuripat and colleagues (2017) found a dominance of attentional gain modulation on sensory activity early in training, which was abolished at late phase of training. This reported change of gain modulation (i.e., early vs. late phase of training) may correspond to our observation of changes in pattern difference between two features (i.e., before vs. after training) in V1. In particular, the lack of gain modulation after extensive training may correspond to the absence of decodable WM content. The inability to decode memorized features, however, does not necessarily mean the absence of information. Instead of a gain mechanism, training was proposed to improve performance via noise reduction (Itthipuripat et al., 2017), to which the MVPA of fMRI data might be insensitive.

An alternative interpretation for the lack of decodable WM information in V1 after training is that training induced synaptic changes in WM storage (Christophel et al., 2017; Masse et al., 2020; Mongillo et al., 2008). Previous studies using computational modeling offered a plausible mechanism of activity-silent short-term retention of WM, where the feature-specific information can be retained in the pattern of synaptic weights even in absence of persistent delay activity (Masse et al., 2020; Mongillo et al., 2008). We thus speculate that such effect of synaptic facilitation might be sufficient for mnemonic processing of features after training without relying upon pattern-level differences. Nevertheless, our current study was not sufficient to discriminate between these two possibilities (i.e., noise reduction vs. synaptic changes). Further investigations with advanced neurobiological techniques are needed to clarify this issue.

Another governing assumption in WM research has been that the retention of WM information is supported by the delay period activity in the frontoparietal network (D’Esposito & Postle, 2015; Sreenivasan & D’Esposito, 2019). Here, we also showed elevated delay period activity in IPS, which was not influenced by training. Although the pattern activity during the delay period did not contain feature information (i.e., orientation), as also reported in other studies (Linden et al., 2012; Riggall & Postle, 2012), combined evidence from fMRI decoding and TMS suggest that after training, IPS became engaged for feature-specific mnemonic processing. Such learning-dependent changes in IPS may contribute to the formation of a more stabilized WM representation in higher-order areas. Combined with the findings in V1, we suggest that discrimination training may cause a transition of feature-specific WM coding at different cortical levels for complementary roles (Bettencourt & Xu, 2016; Lorenc et al., 2018). We should note that IPS may not be the unique region that shows robust delay period activity, as prefrontal cortex (PFC) has also been proposed to associate with mnemonic processing (Ester et al., 2015; Miller et al., 1996; Stokes et al., 2013). However, we did not find meaningful results in PFC from fMRI decoding in the current study. This null finding was also reported by prior studies and may be ascribed to the higher order information representation (e.g., task rules, abstract representations of categories) in PFC (Ester et al., 2015; Lee et al., 2013). Future studies may use neurophysiological methods to investigate learning-dependent modification in the WM representation in PFC.

Previous studies have produced mixed results regarding the involvement of V1 and IPS during WM delay, especially between human and non-human primates. Here, we point to the role of training that may explain such discrepancy. While human only need a few trials to familiarize the task, monkeys have to go through extensive training before recording their neural activity, analogous to the measurement of posttraining performance in human learning studies. In this regard, the decodable WM content found in human V1 (Harrison & Tong, 2009; Serences et al., 2009) versus weak (or absent) WM information reported in sensory areas from neurophysiological studies (Mendoza-Halliday et al., 2014; Zaksas & Pasternak, 2006) may reflect the key difference in training. Similarly, the decodable WM information contained in elevated delay period activity from neurophysiological studies (Averbeck & Lee, 2007; Baeg et al., 2003) versus the mixed results of WM decoding in parietal cortex from human neuroimaging studies (Linden et al., 2012; Riggall & Postle, 2012) may also relate to the difference in training. Thus, we provide a plausible account that can reconcile the discrepant findings across species. That is, the neural locus of WM representation depends critically upon training.

Interestingly, a previous study that used feature discrimination task showed a lack of TMS effect when stimulating IPS both before and after training (Chang et al., 2014a). However, in that study, two features were concurrently, rather than sequentially, shown to the participants. This design did not require short-term memory retention of a particular item and thus heavy involvement of IPS was not necessary for performing the task. The contrast between our results and the findings in Chang et al. (2014) actually supported the critical dependence on parietal cortex to afford stable WM representation after training, specifically when the task included mnemonic processing of the stimuli (Postle, 2006; Xu, 2017).

In summary, discrimination training differentially influenced the mnemonic processing of visual features in sensory and higher-order areas. Although the sensory engagement for WM is relatively independent of training, training may alter the coding format of WM content in this region. In contrast, the recruitment of higher-order parietal areas for WM representation depends upon training, potentially contributing to a more stabilized representation along with improved discriminability. The feature-specific WM representation at different cortical areas may serve complementary roles to support learning-related brain plasticity throughout multiple cortical hierarchy (Dosher & Lu, 2017; Maniglia & Seitz, 2018; Watanabe & Sasaki, 2015).

## Acknowledgements

This work was supported by grants from National Key R&D Program of China (2017YFB1002503) and the National Natural Science Foundation of China (31271081, 31230029, 31800910).

## Notes

### Competing Interest Statement

The authors have declared no competing interest.

